# Vaccination has minimal impact on the intrahost diversity of H3N2 influenza viruses

**DOI:** 10.1101/085985

**Authors:** Kari Debbink, John T. McCrone, Joshua G. Petrie, Rachel Truscon, Emileigh Johnson, Emily K. Mantlo, Arnold S. Monto, Adam S. Lauring

## Abstract

While influenza virus diversity and antigenic drift have been well characterized on a global scale, the factors that influence the virus’ rapid evolution within and between human hosts are less clear. Given the modest effectiveness of seasonal vaccination, vaccine-induced antibody responses could serve as a potent selective pressure for novel influenza variants at the individual or community level. We used next generation sequencing of patient-derived viruses from a randomized, placebo-controlled trial of vaccine efficacy to characterize the diversity of influenza A virus and to define the impact of vaccine-induced immunity on within-host populations. Importantly, this study design allowed us to isolate the impact of vaccination while still studying natural infection. We used pre-season hemagglutination inhibition and neuraminidase inhibition titers to quantify vaccine-induced immunity directly and to assess its impact on intrahost populations. We identified 166 cases of H3N2 influenza over 3 seasons and 5119 person-years. We obtained whole genome sequence data for 119 samples and used a stringent and empirically validated analysis pipeline to identify intrahost single nucleotide variants at ≥1% frequency. Phylogenetic analysis of consensus hemagglutinin and neuraminidase sequences showed no stratification by pre-season HAI and NAI titer, respectively. In our study population, we found that the vast majority of intrahost single nucleotide variants were rare and that very few were found in more than one individual. Most samples had fewer than 15 single nucleotide variants across the entire genome, and the level of diversity did not significantly vary with day of sampling, vaccination status, or pre-season antibody titer. Contrary to what has been suggested in experimental systems, our data indicate that seasonal influenza vaccination has little impact on intrahost diversity in natural infection and that vaccine-induced immunity may be only a minor contributor to antigenic drift at local scales.

## Introduction

Despite recommendations for universal influenza vaccination and the ample availability of vaccines in the United States, influenza continues to cause significant morbidity and mortality [1]. This is, in part, a result of the modest effectiveness of current vaccines, so that considerable numbers of vaccine failures occur each year. Within individuals, influenza populations exist as a collection of closely related, and at times antigenically distinct, variants that may exhibit diverse phenotypes [2-6]. Intrahost single nucleotide variants (iSNV) can be transmitted as part of the infecting population [5,7-10] or generated over the course of an infection due to the virus’ low replication fidelity [11,12]. The evolutionary forces that shape the genetic structure of viral populations within hosts and ultimately give rise to novel antigenic variants at the host population level are poorly understood. A clear understanding of the intrahost diversity of influenza virus populations and its impact on influenza virus evolution is central to many questions of direct clinical and public health relevance [13].

Influenza vaccines are considered for reformulation each year to counter the viral antigenic drift that enables escape from the previous year’s vaccine [14]. Annual influenza vaccine effectiveness is 60% on average, and can be much lower during antigenically unmatched years [15,16]. While antigenic drift is monitored annually on a global scale, the source of antigenic variation is ultimately at the level of the individual host. Phylogenetic studies of whole genome sequences from cities and smaller communities have demonstrated that multiple lineages circulate over the course of a single influenza season [2,17], and individual hosts may harbor mixed infections that include antigenically novel variants [3,4]. While human hosts could be preferentially infected with one lineage over another based on pre-infection immune status, the degree to which circulating escape variants contribute to vaccine failure is currently unknown.

Host immune selection is a major driver of influenza virus evolution on the global scale. Both phylogenetic analysis and antigenic cartography have demonstrated that antibodies exert positive selective pressure on the viral hemagglutinin (HA) and neuraminidase (NA) proteins (23). Vaccination and natural influenza infection often lead to partial, or non-sterilizing immunity, and post-vaccination antibody titers are only a moderate predictor of subsequent protection [18,19]. Previous work has demonstrated that sub-neutralizing concentrations of immune sera can promote the generation of antigenic variants, and some have suggested that vaccination can accelerate the process of antigenic drift [20]. A recent study in vaccinated people suggested that novel antigenic variants could be present at low frequencies [21]. Importantly, humans often differ in their prior exposure to influenza viruses and vaccines, and pre-existing immunity may confound such studies [22]. Therefore, the extent to which partial immunity selects for antigenically relevant variants during natural infection in humans is unclear.

By necessity, most of the available data on vaccination and intrahost evolution have come from analyses of HA sequences in large animal models of infection, including horses, pigs, and dogs. These studies have suggested that intrahost populations include a number of somewhat rare single nucleotide variants that increase and decrease in frequency over the course an infection with sporadic fixation events occurring in some animals. The overall impact of vaccination on antigenic diversification was not clear [7,23-25]. Furthermore, some studies found distinct differences in population structure and diversity between experimentally and naturally infected animals, perhaps due to differences in the infecting strain, the size of the inoculum, and background immunity [7,8,24]. These differences are not unique to animal models and are likely to be an issue in extrapolating results from human experimental challenge models to natural infection [26].

Some have suggested that high intrahost diversity reflects increased viral fitness, and mechanisms that alter intrahost diversity may impact evolutionary trajectories [27-29]. If the transmission bottleneck is sufficiently wide, low frequency variants that arise within a host can plausibly be transmitted and spread through host populations [5,10]. Work in animal models and at least one study in humans support the existence of a loose transmission bottleneck [5,8,24]. Bottleneck size appears to be smaller in vaccinated animals [25], and vaccinated pigs and ferrets shed less virus than immunologically naïve animals [24,30]. Understanding how intrahost diversity is generated and maintained and the extent to which host immune status impacts this diversity may be important for defining influenza virus’ larger evolutionary patterns [21,24,25].

Here we used next generation sequencing to define the impact of vaccine-induced immunity on the intrahost diversity of influenza virus during natural infection. We specifically asked: (i) whether influenza viruses in vaccinated individuals represent escape variants, (ii) whether non-sterilizing antibody responses select for novel antigenic variants within hosts, and (iii) the degree to which vaccine-induced immunity impacts the overall diversity of intrahost populations. Because we analyzed influenza populations from individuals enrolled in a randomized, double-blind, placebo-controlled trial of influenza vaccine efficacy [31-33], we were uniquely positioned to define this aspect of immunity to natural infection.

## Methods

### Subjects and specimens

We characterized host-derived influenza populations archived from a randomized, double-blind, placebo-controlled, clinical trial of influenza vaccine efficacy that ran from the 2004-2005 through the 2007-2008 influenza seasons (ClinicalTrials.gov number, NCT00133523, [31-33]). Each year, healthy adults, ages 18-49, were randomized to receive trivalent inactivated influenza vaccine (IIV), live attenuated influenza vaccine (LAIV), or placebo. The study was approved by the Institutional Review Board of the University of Michigan Medical School, and all human subjects provided informed consent. Throat swab specimens were collected from individuals with influenza-like illness within 7 days of onset; residual specimen material was stored in veal infusion broth (VIB) at −80°C. Viral RNA was extracted from 140μl of VIB using the QIAamp viral RNA mini kit (Qiagen 52906), eluted in 50μl buffer, and stored at −80°C. Hemagglutination inhibition (HAI) and Neuraminidase agglutination inhibition (NAI) titers for subjects in this study were previously measured and reported in [18,34].

### Determination of genome copy number

Quantitative reverse transcription polymerase chain reaction (RT-qPCR) was performed on 5µl RNA from each sample using CDC RT-PCR primers InfA Forward, InfA Reverse, and InfA probe, which bind to a portion of the influenza M gene (CDC protocol, 28 April 2009). Each reaction contained 5.4µl nuclease-free water, 0.5µl each primer/probe, 0.5µl SuperScript III RT/Platinum Taq mix (Invitrogen 111732) 12.5µl PCR Master Mix, 0.1µl ROX, 5µl RNA. The PCR master mix was thawed and stored at 4°C, 24 hours before reaction set-up. A standard curve relating copy number to Ct values was generated based on 10-fold dilutions of a control plasmid run in duplicate.

### Illumina library preparation and sequencing

We amplified cDNA corresponding to all 8 genomic segments from 3µl of the viral RNA using the SuperScript III One-Step RT-PCR Platinum Taq HiFi Kit (Invitrogen 12574). Reactions consisted of 0.5µl Superscript III Platinum Taq Mix, 12.5µl 2x reaction buffer, 8µl DEPC water, and 0.2µl of 10µM Uni12/Inf1, 0.3µl of 10µM Uni12/Inf3, and 0.5µl of 10µM Uni13/Inf1 universal influenza A primers [35]. The thermocycler protocol was: 42°C for 60 min then 94°C for 2 min then 5 cycles of 94°C for 30 sec, 44°C for 30 sec, 68°C for 3 min, then 28 cycles of 94°C for 30 sec, 57°C for 30 sec, 68°C for 3 min. Amplification of all 8 segments was confirmed by gel electrophoresis, and 750ng of each cDNA mixture were sheared to an average size of 300 to 400bp using a Covaris S220 focused ultrasonicator. Sequencing libraries were prepared using the NEBNext Ultra DNA library prep kit (NEB E7370L), Agencourt AMPure XP beads (Beckman Coulter A63881), and NEBNext multiplex oligonucleotides for Illumina (NEB E7600S). The final concentration of each barcoded library was determined by Quanti PicoGreen dsDNA quantification (ThermoFisher Scientific), and equal nanomolar concentrations were pooled. Residual primer dimers were removed by gel isolation of a 300-500bp band, which was purified using a GeneJet Gel Extraction Kit (ThermoFisher Scientific). Purified library pools were sequenced on an Illumina HiSeq 2500 with 2×125 nucleotide paired end reads. All raw sequence data have been deposited at the NCBI sequence read archive (BioProject submission ID: SUB1907046)

### Variant detection

Sequencing reads that passed standard Illumina quality control filters were binned by index and aligned to the reference genome using Bowtie [36]. Single nucleotide variants (SNV) were identified and analyzed using DeepSNV [37], which relies on a clonal control to estimate the local error rate within a given sequence context and to identify strand bias in base calling. The clonal control was a library prepared in an identical fashion from 8 plasmids containing the genome for the respective circulating reference strain and sequenced in the same flow cell to control for batch effects. True positive SNV were identified from the raw output tables by applying the following filtering criteria in R: (i) Bonferonni corrected p value <0.01, (ii) average MapQ score on variant reads >30, (iii) average phred score on variant positions >35, (iv) average position of variant call on a read >32 and <94, (v) variant frequency >0.01. We only considered SNV identified in a single RT-PCR reaction and sequencing library for samples with copy number ≥10^5^ genomes/µl transport media or in two separate RT-PCR reactions and sequencing libraries for samples with copy number 10^3^-10^5^ genomes per µl. Our strategy for variant calling is described in [6] and all code can be found at https://github.com/lauringlab/variant_pipeline.

### Phylogenetic analysis

Consensus nucleotide sequences for the HA and NA proteins were aligned using MUSCLE [38]. The best-fit models for nucleotide substitution was identified using jModelTest v2.1.10 [39]. Maximum likelihood phylogenetic trees were generated using RAxML v8 [40] with a GTRGAMMA model, Genbank sequences for vaccine strains as outgroups, and 1000 bootstraps. Trees were visualized and annotated using FigTree (v1.4.2).

### Data analysis and statistics

All statistical analyses were performed using Prism 6 and R. Description of the analysis and annotated code are available at https://github.com/lauringlab/fluvacs_paper. HA structural models were generated and visualized with PyMol.

## Results

### Study subjects and specimens

We utilized influenza A positive samples from a randomized, double-blind, placebo-controlled study of vaccine efficacy that took place during the 2004-2008 influenza seasons at six study sites in Michigan. This trial measured vaccine efficacy of both the trivalent inactivated (IIV) and live attenuated influenza vaccine (LAIV) compared to placebo and each other. We sequenced patient-derived influenza populations without culturing from three seasons: 2004-2005, 2005-2006, and 2007-2008. The 2006-2007 influenza season did not have enough influenza-positive samples for our study (total n=16). Influenza A (H3N2) strains dominated the other three seasons, and the circulating 2004-2005 virus was considered at the time to be only a modest mismatch with the vaccine strain. The other seasons were antigenically matched. The number of subjects each year was as follows: 2004-2005 season, 522 IIV, 519 LAIV, and 206 placebo [31]; 2005-2006 season, 867 IIV, 853 LAIV, and 338 placebo [32]; 2007-2008 season, 813 IIV, 814 LAIV, and 325 placebo [33]. Over these 5119 person-years of observation, 165 individuals had culture or RT-PCR confirmed influenza A (H3N2) infection and specimens available for analysis. Of these, 80 individuals had received LAIV, 42 had received IIV, and 43 had received placebo. For 2004-2005, flu-positive samples were available for 28 subjects: 7 IIV, 12 LAIV, and 9 placebo, and for 2005-2006, 32 samples were available to study: 13 IIV, 14 LAIV, and 5 placebo. During the 2007-2008 season, 105 flu-positive samples were available: 22 IIV, 54 LAIV, and 29 placebo. We were able to amplify and quantify genomes for 119 of the 165 influenza-positive samples (Table 1). The average age of this sequenced cohort from all years was 24.5, indicating that participants were generally young and likely shared similar influenza pre-exposure histories particularly after randomization. The age, sex and race of the cohort were similar to that of the overall study cohort for each of the 3 seasons.

**Table 1.** Samples analyzed over three FLU-VACS Seasons

|  | 2004-2005 | 2005-2006 | 2007-2008 |
| --- | --- | --- | --- |
| H3N2 Vaccine Strain | Wyoming/3/2003 | California/7/2004 | Wisconsin/67/2005 |
| Circulating Strain(s) | Wyoming/3/2003<br>California/7/2004 | California/7/2004<br>Wisconsin/67/2005 | Wisconsin/67/2005<br>Brisbane/10/2007 |
| Demographics |  |  |  |
| Average Age (Years) | 30.0 | 28.8 | 22.8 |
| Sex (% Female) | 71.4% | 83.3% | 70.1% |
| Race (% White) | 92.9% | 94.4% | 86.2% |
| Sequenced Samples | 14 | 18 | 87 |
| IIV | 4 | 9 | 16 |
| LAIV | 5 | 6 | 44 |
| Placebo | 5 | 3 | 27 |
| Intrahost SNV Data | 10 | 9 | 64 |
| IIV | 3 | 6 | 11 |
| LAIV | 4 | 1 | 30 |
| Placebo | 3 | 2 | 23 |

In the 2007-2008 season, roughly 40% of individuals in the larger cohort reported having ever received a prior influenza vaccination. Despite subject randomization, differences in pre-existing immunity due to prior vaccination or influenza infection could impact results. To control for this possibility, we obtained pre-season antibody titers by hemagglutination inhibition (HAI) and neuraminidase inhibition (NAI) assays for all study participants against that season’s vaccine strain (Supplementary Figure 1). Vaccine-induced antibody titers and overall vaccine efficacy are generally stable for a single flu season [41]. Therefore, our pre-season HAI and NAI titers are likely to be similar to titers at the time of infection. Pre-season (post-vaccination) titers for individuals in the IIV group were above the geometric mean for the entire sequenced cohort, LAIV subjects had titers spanning the mean, and those in the placebo group were generally below the mean for all seasons. These data demonstrate that in the IIV group, and to a lesser degree, the LAIV group, individuals had strain-specific antibody levels sufficient to apply selective pressure against the infecting virus.

**Figure 1.**
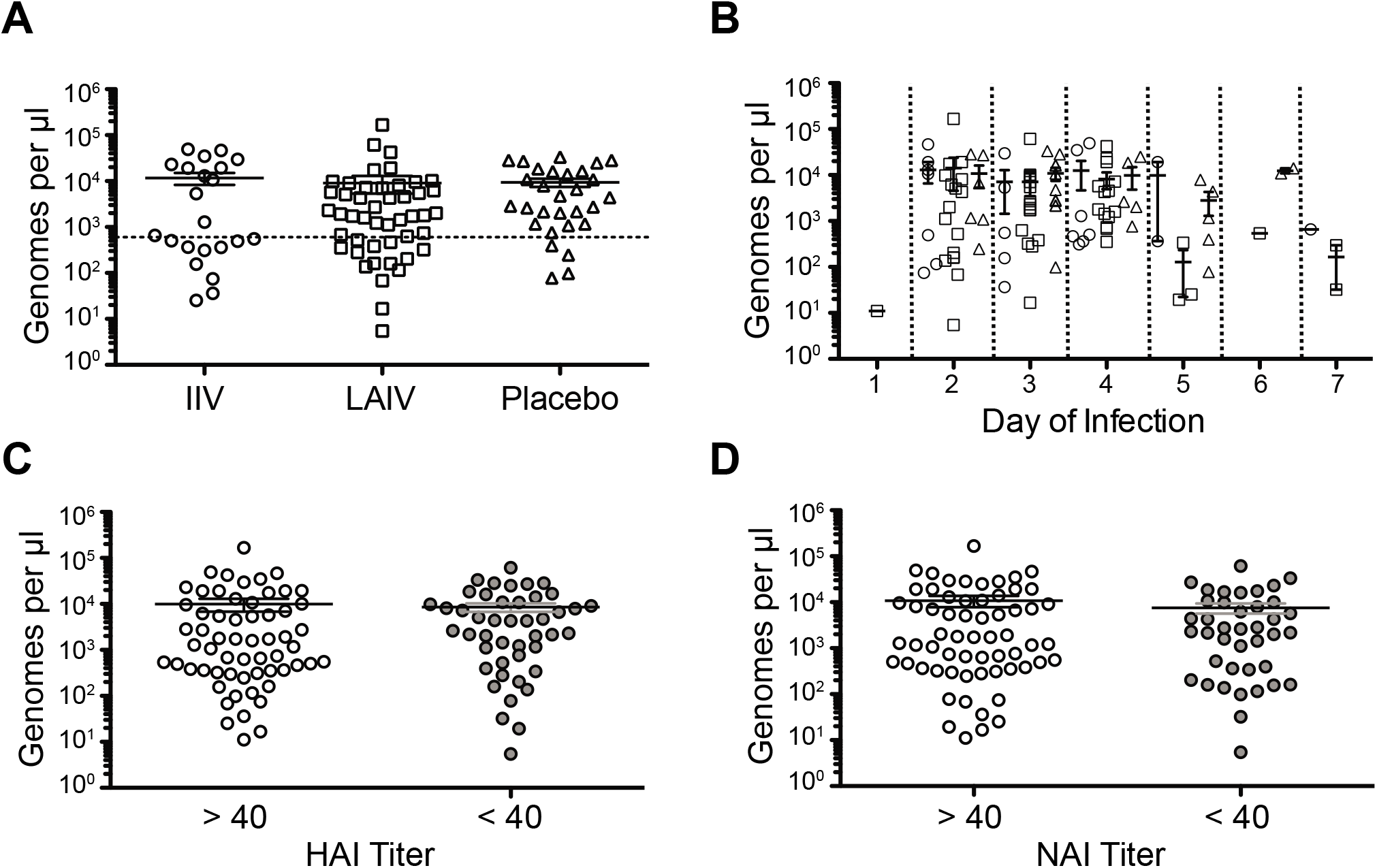
Viral shedding by vaccination status. Genome copy number per µl of transport media was determined by RT-qPCR all samples from the 2007-2008 season. (A) Copy number by vaccination status. IIV, inactivated influenza vaccine; LAIV, live attenuated influenza vaccine. (B) Copy number by day of infection (onset of symptoms is day 0) and vaccination status. Circle, IIV; Square, LAIV; Triangle, Placebo. (C) Copy number by HAI titer (D) Copy number by NAI titer. There were no differences among any of the groups by one-way ANOVA with Bonferroni correction.

### Viral load across groups

We have previously shown that viral load influences the sensitivity and specificity of iSNV detection [6]. In order to determine whether viral load was different among the IIV, LAIV, and placebo samples that we sequenced, we measured genome copy number by RT-qPCR for the 2004-2005, 2005-2006, and 2007-2008 seasons. For the 2007-2008 season, which had the most samples, there were no significant differences in copy number by vaccination group (Figure 1A). In agreement with the 2007-2008 data, we did not detect differences in copy number by vaccination group for the 2004-2005 and 2005-2006 seasons (Supplementary Figure 2). Since copy number is dependent on time from illness onset [42,43], we analyzed the data based on sample collection day (Figure 1B). Using days 2-4, for which there were at least 5 data points for each treatment group, we did not find any significant differences (p=0.24-0.57 for days 2-4, non-parametric one way ANOVA). We divided the larger group of 2007-2008 subjects into groups based on pre-season HAI and NAI titers ≥40 or <40 against that season’s strain, as an HAI titer of 40 is typically considered to be associated with 50% protection given exposure [18,19,44]. This cutoff was identical to the HAI and NAI geometric mean titers for our sequenced cohort (61.9 and 34.5, respectively), given the dilutions used. We did not detect differences in copy number based on HAI or NAI titer (Figure 1C-D), even when accounting for day of symptom onset (p=0.25 for HAI, p=0.97 for NAI, Mann-Whitney U test). Because we only measured copy number in the subset of virus populations that were amplified and sequenced, these data should not be interpreted in the context of vaccination and overall shedding [45].

**Figure 2.**
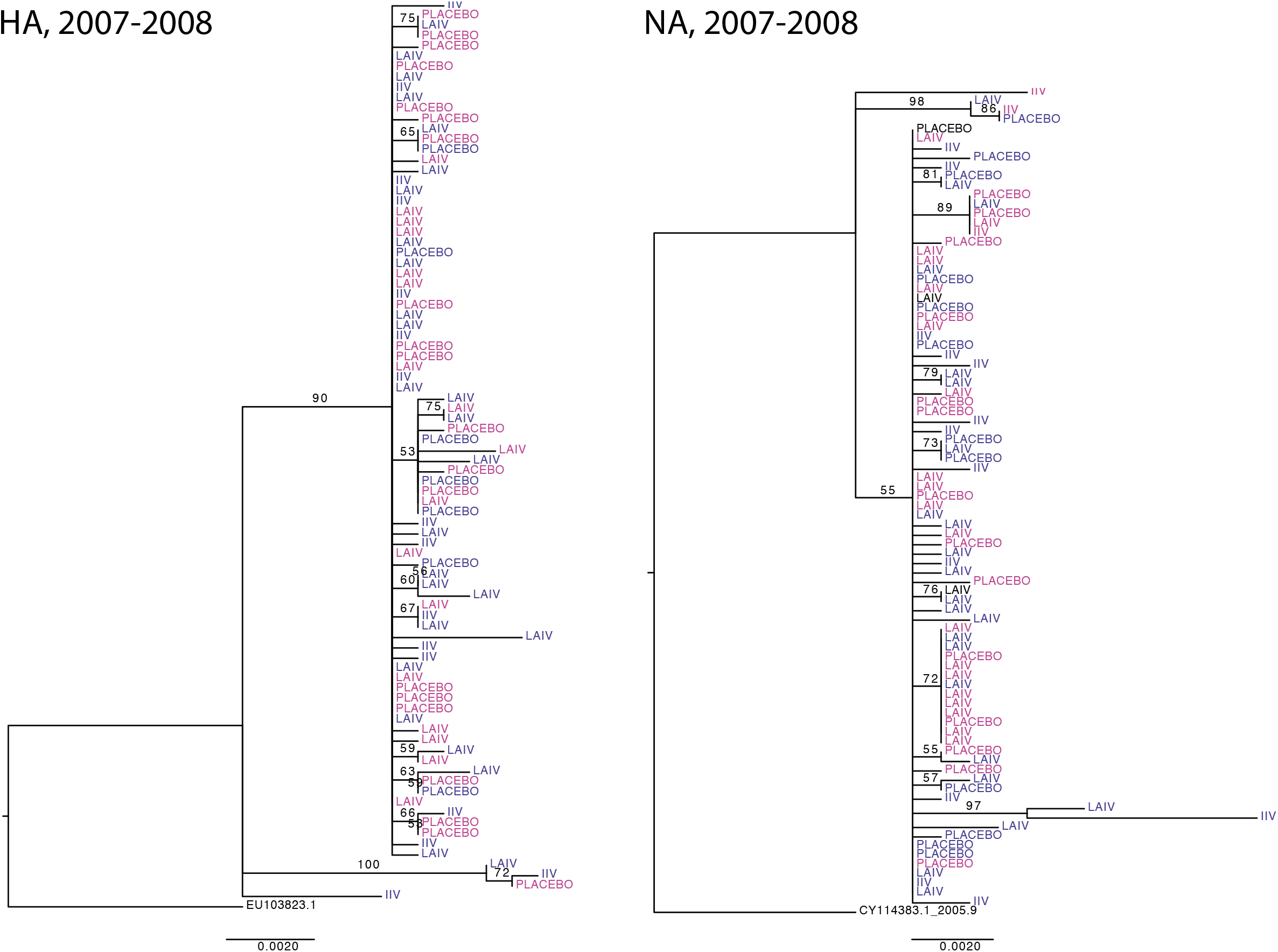
Phylogenetic trees of HA and NA consensus sequences from the 2007-2008 season. Maximum likelihood trees of HA (left) and NA (right) with tips coded by vaccine status and preseason HAI (left; blue >40, magenta <40) or NAI (right; blue >40, magenta <40) titer. Outgroups are HA (EU103823.1) and NA (CY114383.1) for the vaccine strain A/Wisconsin/67/2005. Bootstrap values (n=1000 bootstraps) are shown and nodes with bootstrap values <50 are collapsed for easier visualization.

### Deep sequencing of intrahost influenza populations

We used the Illumina platform to determine the whole genome consensus sequence and to identify intrahost single nucleotide variants for each patient-derived sample. Importantly, we have developed and rigorously benchmarked a variant calling pipeline that maintains high sensitivity for rare iSNV detection while dramatically reducing false positive variant calls [6] (Supplementary Table 1). We have found that the number of false positive iSNV calls is much higher in samples with genome copy numbers <10^3^ per µl of transport media; therefore, we only report iSNV from a high quality dataset that includes 64 samples from the 2007-2008 season. High quality iSNV data for the 2004-2005 and 2005-2006 influenza seasons are included in Supplementary Figures 4, 6, and Supplementary Table 2. Our libraries yielded an average coverage above 20,000 reads per base with even coverage across the coding region of all segments for each season (Supplementary Figure 3).

**Figure 3.**
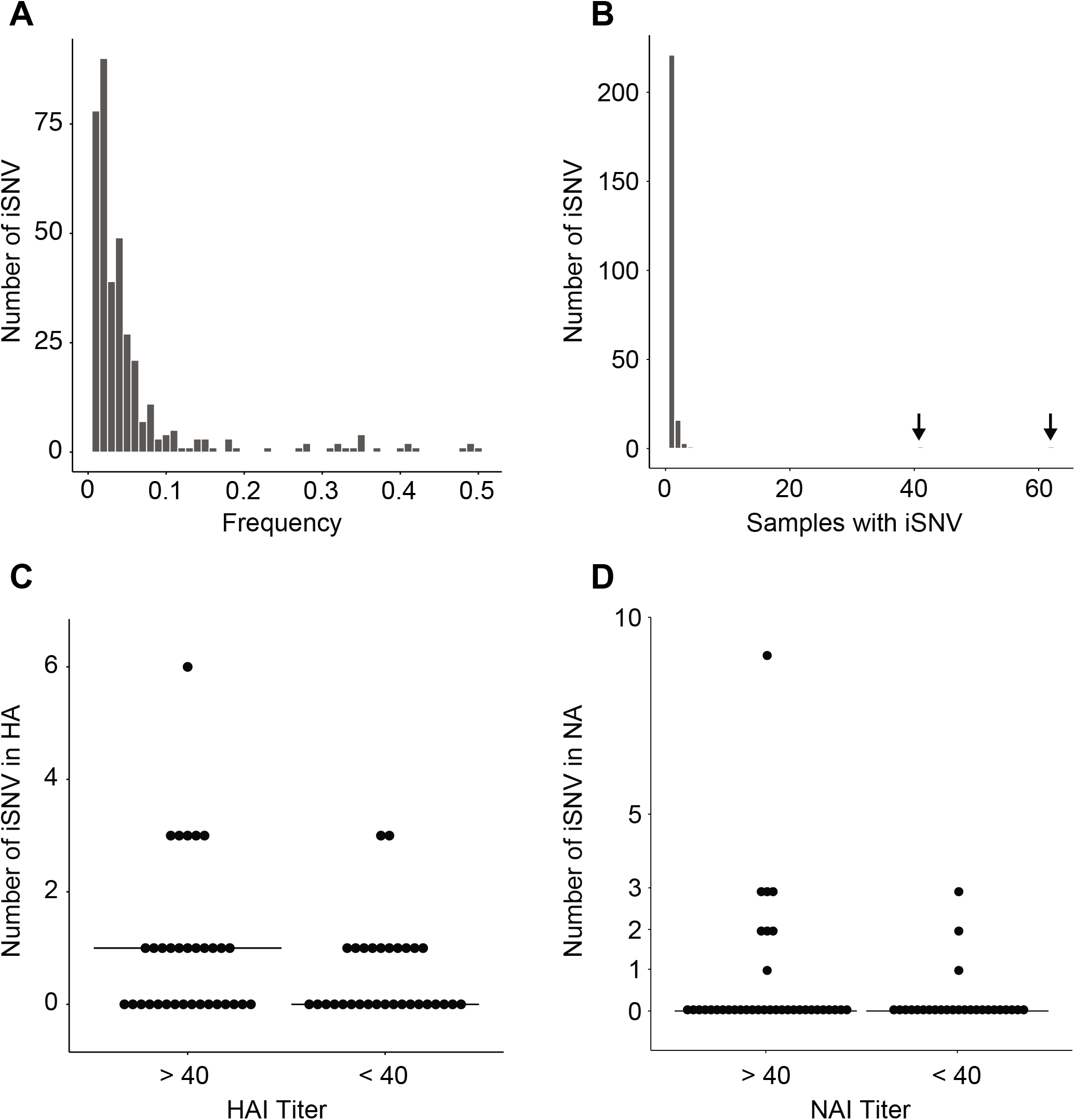
Intrahost diversity in samples from the 2007-2008 season. (A) Histogram of the number of iSNV at a given frequency. Bin width = 0.01. (B) Histogram of the number of samples in which each iSNV is found. Arrows indicate bars with one SNV, which are hard to discern in the histogram. These polymorphic SNV, at PB2 position 900 and PA position 515 respectively, were found at 4-6% frequency within hosts and in similar numbers of individuals across vaccination groups. (C) Number of HA iSNV per sample stratified by pre-season HAI titer. >40 = serologically immune, <40 = not serologically immune. (D) Number of NA iSNV per sample stratified by pre-season NAI titer. >40 = serologically immune, <40 = not serologically immune.

**Table 2.** Number of iSNV (mean ± interquartile range) by segment and treatment group for samples from the 2007-2008 season.

| Segment | IIV (n=11) | LAIV (n=30) | Placebo (n=23) |
| --- | --- | --- | --- |
| 1 (PB2) | 2.1 ± 2 | 1.78 ± 1 | 1.89 ± 1 |
| 2 (PB1) | 1.8 ± 1 | 1.46 ± 1 | 1.45 ± 1 |
| 3 (PA) | 1.8 ± 0.75 | 1.7 ± 1 | 1.65 ± 1 |
| 4 (HA) | 1.43 ± 1 | 1.27 ± 0.5 | 1.27 ± 0 |
| 5 (NP) | 1.75 ± 1.25 | 1.33 ± 0.75 | 1.4 ± 1 |
| 6 (NA) | 1 ± 0 | 1.17 ± 0 | 1.25 ± 0.25 |
| 7 (M) | 1 ± 0 | 1.14 ± 0 | 1 ± 0 |
| 8 (NS) | 1 ± 0 | 1.33 ± 0.75 | 1.29 ± 0.5 |

### HA and NA sequences do not cluster by vaccination status or pre-season antibody titer

Given the frequent observation of community-level diversity in circulating influenza viruses [2,17], we asked whether vaccinated individuals were infected with distinct strains relative to the placebo group. If vaccine failures were due to infection with an antigenically distinct variant, we would expect to see evidence of clustering by vaccination- or sero-status in HA and NA phylogenetic trees. We therefore analyzed HA and NA consensus sequences from all 87 individuals in the 2007-2008 season (Figure 2). There was very little diversity in either gene, and we found that sequences from individuals in each treatment group (e.g. IIV, LAIV, or placebo) were dispersed throughout the tree. More importantly, we found no evidence for clustering based on pre-season HAI and NAI titer (by colors in Figure 2). We obtained similar results from the 2004-2005 and 2005-2006 seasons, albeit with fewer sequenced samples (Supplementary Figure 4). These data suggest that within-season and within-host antigenic drift due to high levels of vaccine-induced antibodies are not major determinants of vaccine failure.

**Figure 4.**
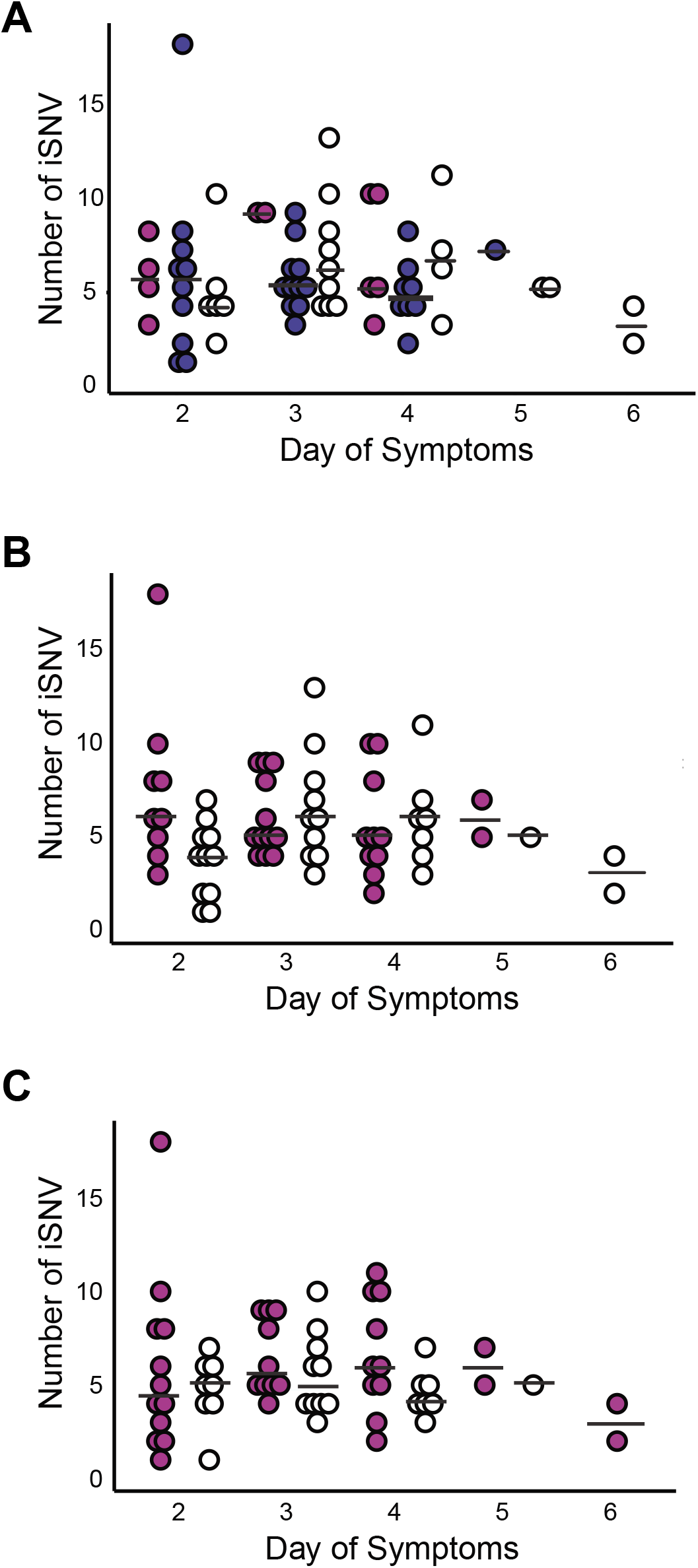
Temporal patterns of intrahost diversity. Number of genome-wide iSNV per sample (y-axis) by day of symptoms (x-axis) stratified by (A) recipients of IIV, magenta; LAIV, blue; placebo, white (B) HAI >40, magenta; HAI <40, white (C) NAI >40, magenta; NAI <40, white. Mean number of iSNV in each group is indicated (bar).

### Intrahost diversity in vaccinated and unvaccinated individuals

We next analyzed the iSNVs present in 64 of our samples from the 2007-2008 season. We identified 369 minority variants across the entire genome, most of which were present at a frequency of <0.1 (Figure 3A). The vast majority of iSNV were only found once (Figure 3B). We also evaluated whether partial immunity impacts viral diversity by comparing the number of iSNVs per sample in the HA and NA genes based on HAI and NAI titers, respectively. We did not observe a difference in iSNV count based on HAI and NAI titers for either HA or NA (Figure 3C-D, p=0.20 for HA, p=0.26 for NA, Mann Whitney U test). The average number of iSNV per sample was similar across the genome regardless of host treatment group (Table 2, p=0.38, non-parametric one way ANOVA) or HAI and NAI titers (Supplementary Figure 5, p=0.12 for HAI, p=0.22 for NAI, Mann Whitney U test).

**Figure 5.**
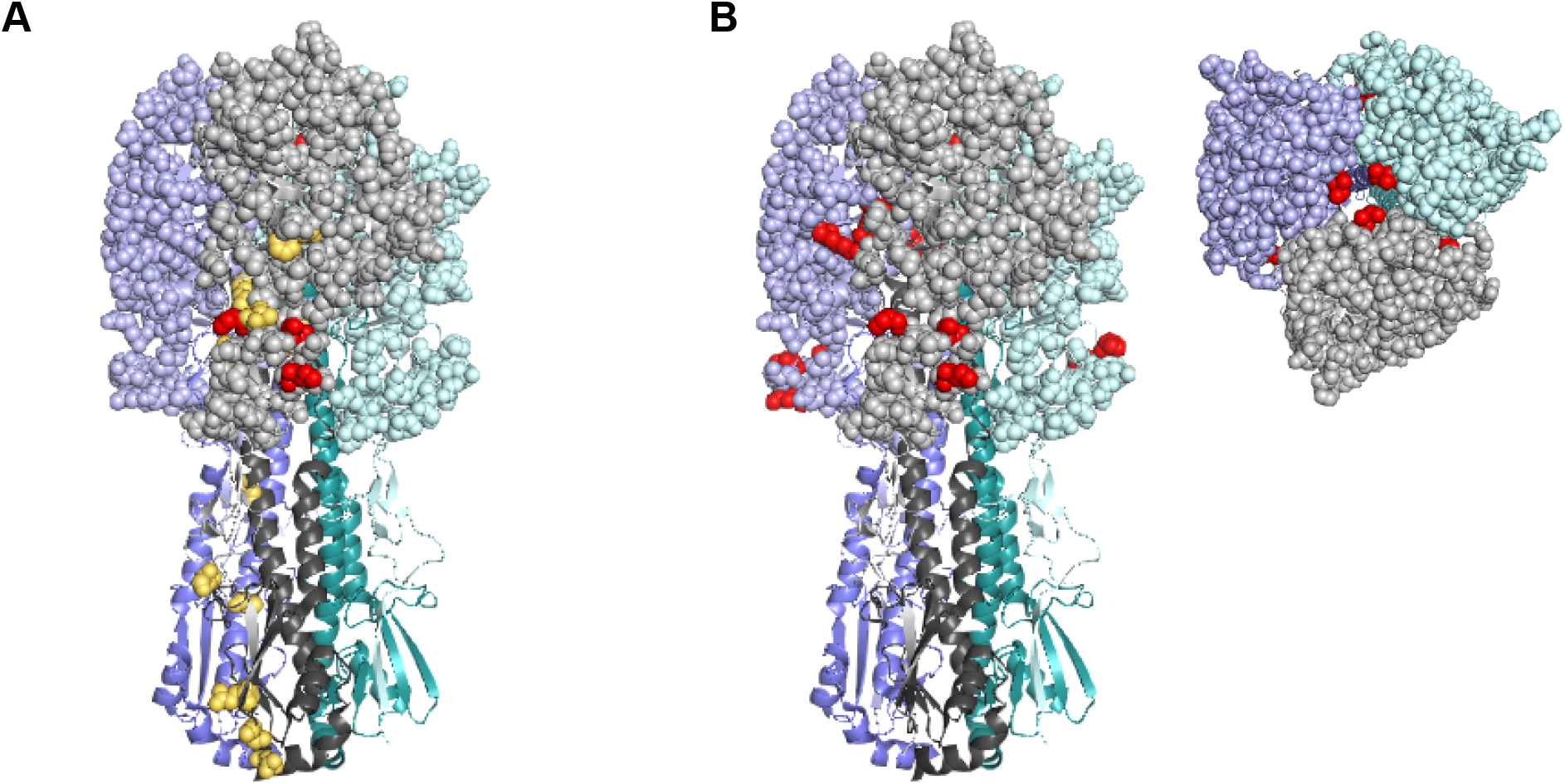
Structural mapping of HA variants. A homology model of the A/Brisbane/10/2007 (H3N2) HA trimer is shown, with each monomer represented by a different color (purple, grey, and teal) and HA1 and HA2 designated by lighter and darker shades of the same color, respectively. (A) A side view of HA. All identified non-synonymous mutations and known antigenic amino acid positions are shown as balls on the grey monomer. Those variants colored red are within known antigenic sites, while light orange mutations are not. (B) A side and top view of HA. All amino acid positions within known antigenic sites are displayed as balls, with the antigenic mutations identified here shown in red on all three monomers.

Most iSNV from the 2004-2005 and 2005-2006 seasons were also found in only one sample each and at frequencies <0.1 (Supplementary. The number of iSNV did not differ based on treatment group or HAI and NAI titers (Supplementary Table 2). We did not identify any variants specifically associated with vaccination group or HAI and NAI titers for any of the influenza seasons analyzed. Unlike experimental challenge studies, we only had one sample per person and could not evaluate changes in diversity at the level of the individual host. However, we did not identify any significant differences in diversity by day of infection across the cohort by vaccination status (Figure 4, p=0.15-0.79 for days 2-4, non-parametric one way ANOVA) or antibody titer. We identified only a marginal difference in the number iSNV across the genome based on HAI titer on day 2 (uncorrected p-value 0.02 with >6 comparisons, Mann Whitney U test). These data suggest that our results are unlikely to be confounded by temporal sampling issues.

### Evidence for low antigenic diversity within hosts

Some have suggested that vaccine-induced immunity will select for novel antigenic variants within hosts [21]. We therefore compared iSNV in the HA gene by vaccination group and serostatus. Of the 17 variants we identified that resulted in nonsynonymous changes within HA, 11 were in HA1 and 6 were in HA2 (Figure 5, Supplementary Table 3). Five antigenic sites, comprising 131 amino acid positions, have been described in HA1 for H3N2 viruses [46-48]. Six of the HA1 variants identified in our study were located in antigenic sites C and D (Figure 5B), one of which was found in two samples (Supplementary Table 3). Of these potential antigenic variants, three were found in vaccinated individuals (1 IIV, 2 LAIV) and four were found in those in the placebo group. When grouped by HAI titer, four were found in samples from individuals with titers ≥40, while three were found in individuals with titers <40. No variants were specific to vaccinated individuals, and antigenic diversity was similar across all groups. None of the identified mutations were observed in subsequent circulating H3N2 strains.

## Discussion

We set out to define the relationship between vaccine-induced immunity and the intrahost diversity of influenza virus. We hypothesized that non-sterilizing immunity could potentially select for novel antigenic variants and contribute to larger scale patterns of influenza evolution. We were able to definitively address these questions using samples derived from a randomized, placebo-controlled vaccine trial in healthy, young adults who likely had similar prior exposure to influenza viruses [31,33]. The availability of post-vaccination, preseason HAI and NAI titers allowed us to examine directly the impact of measured serologic antibody pressure rather than using vaccination alone as a surrogate marker. Because all individuals were infected naturally, our data provide a rare view of within-host influenza virus diversity in humans. We directly sequenced the samples without passage in cell culture, eliminating the possibility of culture-adapted mutations and employed a well-benchmarked variant calling pipeline [6] that dramatically reduces the false positive iSNV calls that often plague next generation sequencing studies. In this exhaustive and well-controlled study, we found no differences in intrahost influenza diversity based on vaccination status or HAI and NAI titers.

Our findings in a natural infection system are concordant with an equine influenza virus evolution study in vaccinated horses. Intrahost variation was similar between naïve and vaccinated horses, regardless of whether they were infected naturally or experimentally [25]. However, not all experimental infection studies mirror results seen during natural infection. A study investigating swine influenza virus found discrepancies in intrahost variation based on whether animals encountered natural infection or were experimentally infected [24]. Our data is in contrast with a study of experimentally-infected dogs that uncovered differences in intrahost diversity and evolution in antigenic sites based on vaccination status [23]. Two other studies of equine influenza virus found mixed infections of multiple influenza lineages during natural infection, which would not be seen in an experimental model but may be relevant to the transmission and spread of novel variants [7,8].

We did not detect phylogenetic clustering of HA and NA sequences based on vaccination status, type of administered vaccine, or pre-season HAI and NAI titers. These results are consistent with those of Dinis et al., who found no segregation of HA sequences based on vaccination status in a case test-negative study of vaccine effectiveness [21]. Together, these data suggest that vaccinated and unvaccinated individuals are infected with similar strains and that within season antigenic drift is not a major contributor to reduced vaccine efficacy. Because we also stratified our analysis by pre-season HAI and NAI titer, we can similarly exclude viral escape from a non-sterilizing antibody response. Our data further suggest that pre-existing antibody against circulating strains does not apply sufficiently strong selective pressure to drive the emergence of antigenically distinct strains within a given host.

High intrahost diversity may be an important factor in viral evolution, since it increases the number of novel variants on which natural selection can act. Some have proposed that the intrahost diversity of RNA viruses is linked to virulence [27,28], suggesting that processes that act to restrict or enhance intrahost diversity may alter disease phenotypes. We found that within host diversity of influenza virus was quite low. Most iSNV were present at frequencies of less than 0.1, which means they would only plausibly be spread between hosts if the transmission bottleneck were reasonably large [5]. The number of iSNV in a given host was similar between vaccine and placebo groups, both across the genome and on the segments coding for HA and NA. Furthermore, there were no significant differences in diversity when samples were grouped by HAI and NAI titer, and we did not find evidence of treatment-specific iSNV. Together our data suggest that vaccine-induced immunity does not significantly influence intrahost diversity and is a relatively weak selective pressure at the level of the individual host.

While the number of iSNV was similar in all groups, we considered it possible that partial immunity may drive the emergence of specific antigenic variants. Vaccination and natural infection can induce a wide range of immune responses depending on host and viral factors. These can range from complete protection to leaky responses that allow infection, but influence disease severity or duration. Non-sterilizing immune responses have the potential to select for escape variants within each host. In the setting of current recommendations for universal vaccination, a highly vaccinated population could potentially select for antigenically evolved viruses more quickly than the spread of natural infection. For this reason, it is important to tease apart the role of vaccination on influenza evolution. We did find a number of variants in antigenic sites, but these were no more frequent than in other regions of the genome and did not vary with vaccination status or HAI and NAI titer. Importantly, our results are in conflict with earlier work in several model systems that demonstrate the rapid selection of antigenic variants in the presence of sub-neutralizing antibody or in experimentally infected animals [24,25,49-51]. This discrepancy may be due to differences in infectious dose, host genetic background, history of prior influenza infection, immune correlates not captured by HAI and NAI titers, or the strains tested. Animal studies are often performed in immunologically naïve or genetically identical animals, whereas humans have complex genetic backgrounds and immunological histories that could play a role in mediating population-wide immunity.

While global patterns of influenza transmission and evolution are complex, our study provides important insights regarding influenza evolution on intra- and interhost scales. We were able to define aspects of intrahost evolution within a geographically-constrained and relatively young cohort with similar vaccination histories and previous influenza exposures across groups, limiting potential confounding factors. Still, we acknowledge that by reducing these confounders, we may be missing important determinants of intrahost evolution. For example, intrahost diversity may be different in children, older adults, or those with high-risk conditions. Our study included 5119 person-years of observations to yield a dataset of 165 viruses, 119 of which were sequenced. Large, placebo-controlled, randomized influenza vaccine trials involving thousands of people are unlikely to be conducted in the future due to the recommendation for universal vaccination. Therefore, our sample set likely represents the best chance to directly assess the impact of vaccination on influenza evolution in the context of natural human infection.

Despite our well-controlled study, we found little evidence for vaccine-driven evolution in the context of the community that was sampled. None of the low frequency variants identified in the antigenic sites were found in subsequent seasons. While we cannot rule out the possibility that evolutionary patterns would be different in other geographic regions, different influenza seasons, or with other subtypes, we did not uncover differences between vaccinated and unvaccinated populations with respect to genome-wide or antigenic diversity in three H3N2 seasons. We were not able to evaluate the impact of vaccination on H1N1 or subtype B viruses. Furthermore, our sample set cannot discern whether there are differences in evolutionary pressure based on vaccine match/mismatch, as the vast majority of our samples were from antigenically matched seasons. A major limitation of our study was that only one sample was available for each participant, so we could not track changes in diversity or mutation accumulation within each individual over the course of their infection. However, we did not observe significant differences in the number of mutations over the first 6 days of infection, which is consistent with previous work in horses [25] and a deep sequencing study of 7 humans in an experimental challenge model [52]. Together with these works, our data suggest that within host dynamics are dominated by purifying selection with the transient appearance of minority variants and little sustained fixation. Selection and transmission of antigenically and epidemiologically important variants is likely to be a rare event when studied at this scale. Given that our data are derived from 165 incident infections in 5119 season-years of observation, detection of such events in the course of a natural infection would require an unrealistic sample size.

We evaluated the potential for vaccine-induced immunity to drive intrahost evolution, as this is an important issue in light of the current recommendation for universal influenza vaccination. We did not find evidence for vaccine-induced pressure on the intrahost or consensus level, despite employing several methods used to study evolutionary processes. Our study is larger than other reports and involves extensive analysis of both placebo and vaccination groups. While randomization, placebo control, and reliance on a young healthy population allowed us to address this question in a rigorous manner, these factors may have lessened person-to-person variation. By better defining the intrahost evolutionary mechanisms and how they impact population-wide influenza evolution, we hope to address questions of import to clinicians and public health workers, improve vaccine design, and develop more efficient epidemiological control measures.

